# Retinal stem cells modulate proliferative parameters to coordinate post-embryonic morphogenesis in the eye of fish

**DOI:** 10.1101/437269

**Authors:** Erika Tsingos, Burkhard Höckendorf, Thomas Sütterlin, Stephan Kirchmaier, Niels Grabe, Lázaro Centanin, Joachim Wittbrodt

## Abstract

A fundamental question in biology is how anatomically and functionally distinct tissues coordinate to direct growth and shape in complex organs. We address this question using as a model the eye of teleost fish, which grow while maintaining the precise shape needed for vision throughout the animal’s life.

Combining clonal analysis in the eye of the teleost medaka (*Oryzias latipes*) with a computational agent based model, we find that the neural retina (NR) and retinal pigmented epithelium (RPE) differentially modulate cell divisions to coordinate their growth rates. Cell divisions in the NR are less stochastic, consistent with an upstream role as an inducer of growth in nearby tissues. Cells in the RPE display much higher stochasticity, consistent with a downstream role responding to inductive signals.

Our simulation predicts that the segregation of stem- and progenitor cell domains in the retinal ciliary marginal zone niche is an emergent property, as the topology of the niche preconditions the system to undergo a spatially biased stochastic neutral drift. Clone properties in the NR support this prediction, and further suggest that NR cells control the direction of division axes to regulate organ shape and retinal cell topology.

This work highlights an as yet unappreciated mechanism for growth coordination in a complex organ, where one tissue integrates external and internal cues as a hub to synchronize growth rates in nearby tissues. In the eye of fish, proliferation parameters of neuroretinal stem cells are a minimal target node for evolution to exploit to adapt whole-organ morphogenesis in a complex vertebrate organ.

## Introduction

Teleost fish grow throughout their lives, increasing massively in size (Johns and Easter, 1977). The teleost medaka (*Oryzias latipes*) grows roughly ten-fold from hatching to sexual maturity within 2-3 months (Figure 1 Supplement 1 A). Unlike embryonic morphogenesis, during post-embryonic growth all organs must scale with the increasing body size while fully functioning. In the eye, continuous growth must be additionally balanced with continuous shape-keeping: Proper optics, and thus vision, requires a precise 3D shape. Highly visual shallow water fish such as medaka have near-perfect hemispherical eyes (Fernald, 1990; Nishiwaki *et al.*, 1997; Beck *et al.*, 2004). Thus, the eye of fish provides an excellent system to explore how anatomically and functionally distinct tissues coordinate to grow and maintain the shape of an organ in functional homeostasis (Johns and Easter, 1977; Centanin *et al.*, 2014).

**Figure 1.**
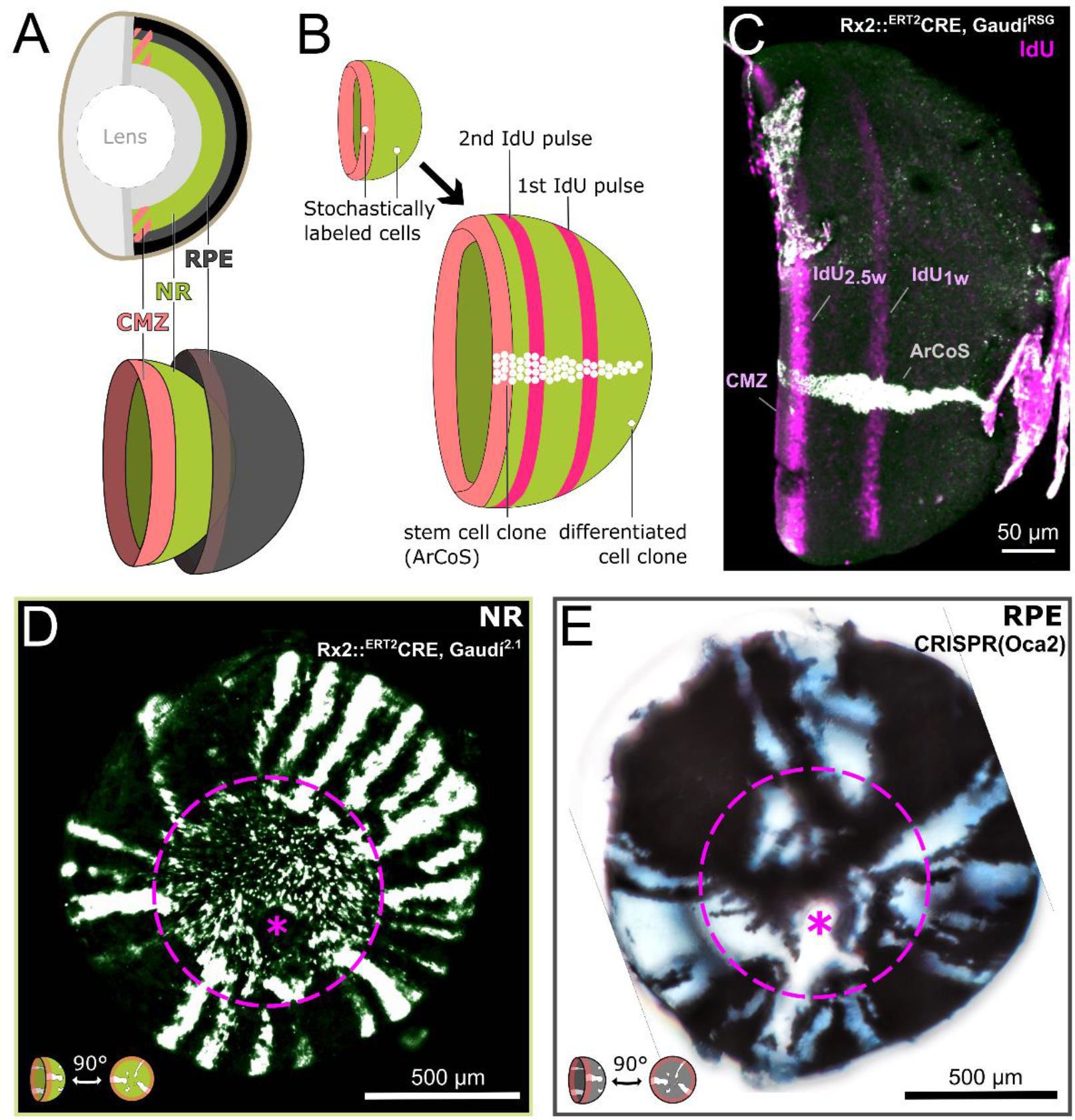
Clonal growth pattern differs in NR and RPE despite identical topology. (**A**) Schematic anatomy of the fish eye. The ciliary marginal zone (CMZ) is a ring-shaped niche that contains independent stem cells for neural retina (NR) and retinal pigmented epithelium (RPE). (**B**) Growth patterns of retinal cell population (concentric rings) and individual clones. Stem cell clones are termed Arched Continuous Stripes (ArCoS). (**C**) False color immunostained NR of 3-week old Rx2::^ERT2^Cre, Gaudí^RSG^ fish with ArCoS and concentric rings of IdU-labelled cells. Overnight IdU pulses were at 1 and 2.5 weeks (w) of age. Leftover undissected autofluorescent tissue fragments cover the far right of the cup-shaped retina. (**D**) Proximal view of clones induced in the NR of Rx2::^ERT2^Cre, Gaudí^2.1^ fish. Maximum projection of confocal stack of GFP immunostaining in false colors; rotated to place optic nerve exit (pink asterisk) in ventral side. Embryonic retina circled with pink dashed line. (**E**) Proximal view of unpigmented lineages induced in the RPE by mosaic biallelic knockout of Oca2 using CRISPR/Cas9. Focused projection of brightfield focal stack; rotated to place optic nerve exit (pink asterisk) in ventral side. Embryonic retina circled with pink dashed line.

The vertebrate eye consists of multiple concentric tissues, including the neural retina (NR) and the retinal pigmented epithelium (RPE) (Figure 1 A). In fish and amphibians, these tissues grow from a ring-shaped stem cell niche in the retinal periphery: the ciliary marginal zone (CMZ) (Johns, 1977; Harris and Perron, 1998; Amato *et al.*, 2004). The CMZ can be subdivided into a peripheral stem- and a central progenitor cell domain (Raymond *et al.*, 2006; Centanin *et al.*, 2014; Wan *et al.*, 2016; Shi *et al.*, 2017). At the very periphery of the CMZ, about 5 rows of cells express the stem cell marker retina-specific homeobox gene 2 (Rx2) (Reinhardt *et al.*, 2015; Wan *et al.*, 2016; Tang *et al.*, 2017). The CMZ is a bi-partite niche, with tissue-specific stem cells for NR and RPE (Shi *et al.*, 2017). In medaka, stem cells for NR and RPE are strictly separate, as demonstrated by transplantations at blastula stage and genetic recombination after hatching (Centanin *et al.*, 2011, 2014). Thus, medaka NR and RPE are independently growing tissues with identical topology.

As a population, CMZ cells appositionally add new cells in concentric rings as shown by label incorporation with thymidine analogues (Johns, 1977; Centanin *et al.*, 2011). Individual stem cells labeled by genetic markers form clonal progeny in so-called Arched Continuous Stripes (ArCoS; Figure 1 B) (Centanin *et al.*, 2011, 2014). Medaka NR stem cells produce the full complement of neuronal cells in apico-basal clonal columns (Centanin *et al.*, 2011, 2014; Lust and Wittbrodt, 2018). These differentiated retinal cells grow little in size (Johns, 1977), retain their relative position over time (Johns, 1977; Centanin *et al.*, 2011), and have negligible death rates (Johns and Easter, 1977; Stenkamp, 2007). Thus, the only parameter available to NR and RPE to coordinate their growth rates is the proliferation of the tissue-specific CMZ stem cells.

Stem cells have long been defined by an unlimited self-renewal capacity (Watt and Hogan, 2000; Clevers and Watt, 2018). Two general strategies underlie long-term maintenance of stem cells: 1) a highly deterministic model where every single stem cell division produces a stem- and a progenitor daughter cell “invariant asymmetry”); and 2) a stochastic model where stem cells divide symmetrically, and the daughter cells have a finite probability to stay as stem cells or commit to a progenitor fate “neutral drift”) (Watt and Hogan, 2000; Clevers and Watt, 2018). One tenet of this model is that stem cells undergo neutral competition: Cells randomly displace one another, resulting in the stochastic “loss” of lineages where all progeny commit to a progenitor fate until the entire niche is occupied by a single clone (Colom and Jones, 2016; Clevers and Watt, 2018).

Strikingly, the medaka retina diverges from the neutral drift model. The CMZ maintains a polyclonal stem cell population for both the NR and the RPE, and in particular NR stem cells undergo asymmetric self-renewing divisions throughout the life of the animal (Centanin *et al.*, 2011, 2014). It remains unclear whether stem cell proliferation in the CMZ follows a purely deterministic model, or whether it poses a new paradigm in-between invariant asymmetry and neutral drift.

In this work we combine *in vivo* and *in silico* clonal analysis in the NR and RPE of medaka to address how these tissues coordinate their growth rates. We find that RPE stem cells divide highly stochastically consistent with a downstream role in the control hierarchy, whereas NR stem cells display less stochasticity consistent with an upstream role in inducing growth in nearby tissues. Our simulation predicts that the spatial segregation of stem and progenitor CMZ domains is an emergent property, as the topology of the retinal niche preconditions the retina to a spatially biased neutral drift. NR stem cells deviate from a purely random drift model by preferential division axis orientation and differential modulation of division parameters along the CMZ circumference. We propose that during post-embryonic growth of the teleost eye, the NR CMZ forms a hub for integrating external and internal stimuli that affect cell division parameters, which ultimately direct the growth and shape of the entire eye.

## Results

### NR and RPE: same topology but different post-embryonic clonal growth modes

The growth rates of all eye tissues must perfectly match, otherwise the organ would deform, akin to a bimetallic strip. The neural retina (NR), and the retinal pigmented epithelium (RPE) are concentric tissues growing from independent stem cells in the ciliary marginal zone (CMZ) (Figure 1 A). Retinal cells follow an exquisite spatiotemporal order (Figure 1 B-C, Figure 1 Supplement 1 B). Thus, clones derived from stem cells (Arched Continuous Stripes; ArCoS) are a frozen record of past cell divisions (Centanin *et al.*, 2011, 2014).

We experimentally generated NR ArCoS by stochastically labelling individual NR stem cells using the Rx2::^ERT2^Cre, Gaudí^2.1^ line in hatchling medaka, and analyzing the eyes in adult fish as previously described (Centanin *et al.*, 2014; Reinhardt *et al.*, 2015). The Rx2 promoter drives the inducible Cre recombinase in stem cells at the very periphery of the CMZ (Reinhardt *et al.*, 2015). A recombined stem cell generates a stripe of GFP-positive progeny in an otherwise GFP-negative retina (Centanin *et al.*, 2014). In proximal view, NR ArCoS emanated as rays from the central embryonic retina, the part of the eye that was already differentiated at the timepoint of Cre-mediated recombination (Figure 1 D).

We visualized RPE ArCoS by mosaic knockout of pigmentation using CRISPR/Cas9 targeted to the gene oculo-cutaneous albinism 2 (Oca2), which is required for melanosome maturation (Fukamachi *et al.*, 2004). RPE stem cells with a bi-allelic mutation in Oca2 generate unpigmented stripes, analogous to RPE ArCoS obtained by transplantation (Centanin *et al.*, 2011). RPE ArCoS frequently branched, forming irregular stripes variable in size and shape (Figure 1 E). These differences in clonal pattern suggested that despite their identical topology, the division behavior of NR and RPE stem cells differed.

### A minimal complexity 3D agent-based model of retinal tissues

To integrate the current knowledge of retinal growth and swiftly address different hypotheses, we built a 3D cell-centered agent based model using the platform EPISIM (Sütterlin *et al.*, 2013, 2017). This modelling technique represents cells as discrete objects (*e.g.* spheres) that physically interact through forces acting on the cell centers; the spheres are allowed to slightly overlap to simulate cell deformability and allow a tight cell packing (Sütterlin *et al.*, 2013, 2017)

Our retinal tissue model consists of a layer of spheres (representing either NR or RPE cells) on a hemisphere (representing the rest of the organ that is not explicitly modelled; Figure 2 A). The RPE is a monolayer, thus each model cell corresponds to one RPE cell. In the NR, CMZ stem cells form a monolayer (Johns, 1977; Raymond *et al.*, 2006), their differentiated progeny forms multi-layered clonal columns (containing all neuroretinal cell types) with little spread tangential to the retinal surface (Figure 1 Supplement 2) (Centanin *et al.*, 2011, 2014; Lust and Wittbrodt, 2018). We took advantage of this fact to not further increase model complexity. Thus, each virtual stem cell represents one NR stem cell, while each differentiated model cell represents a clonal retinal column in the NR.

**Figure 2.**
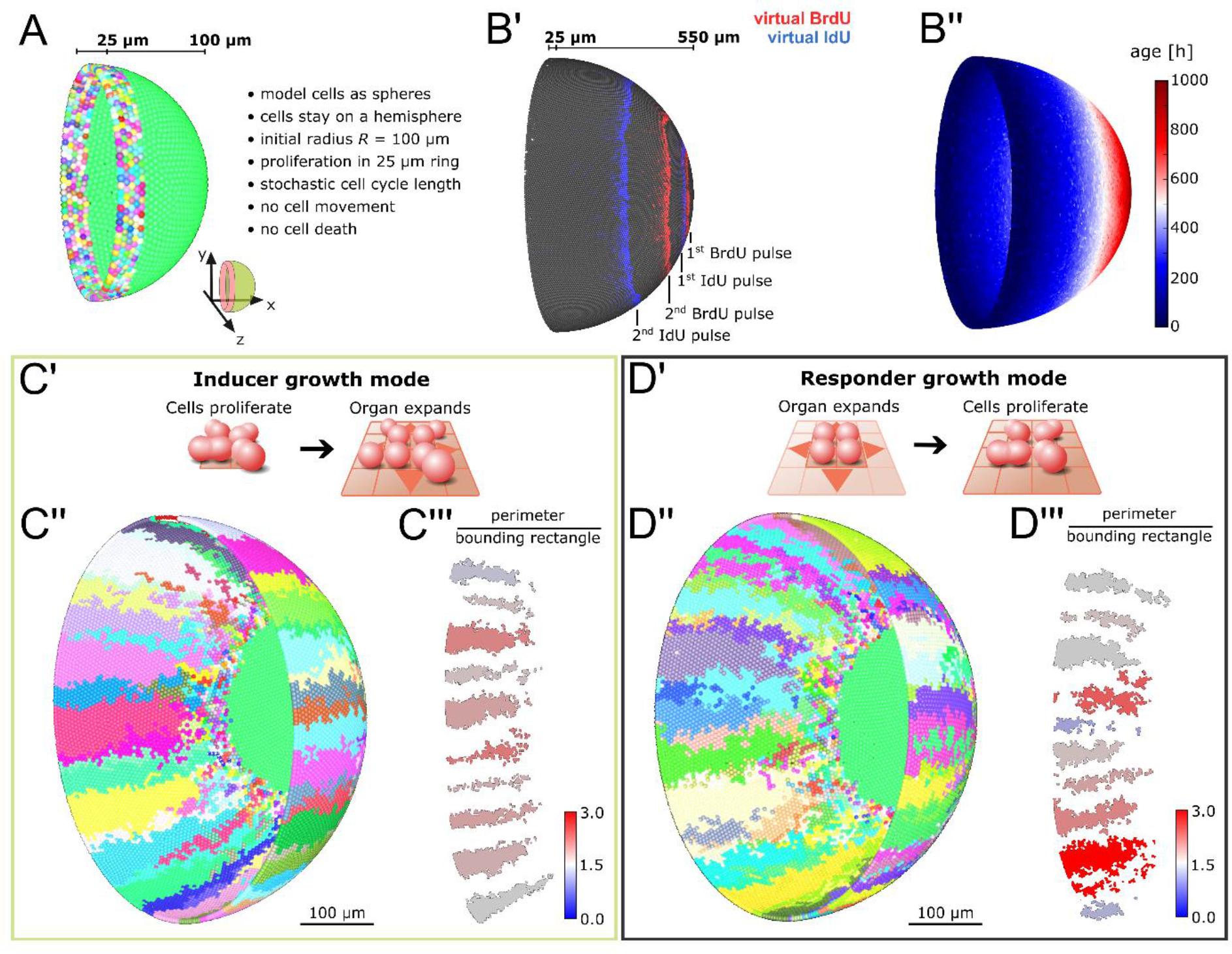
Feedback between proliferation and organ growth affects the simulated clonal pattern. (**A**) Initial condition and properties of the agent-based model of the growing fish retina. Virtual embryonic retina in light green. CMZ cells are assigned unique colors for virtual clonal analysis. (**B’**) Simulated BrdU and IdU pulse-chase experiment. Pulses started at simulation steps 50 and 500 for BrdU, 150 and 1000 for IdU. Each pulse lasted 30 simulation steps, which on average captured one cell cycle. Virtual cells incorporate BrdU/IdU when they divide and half of the signal is passed on to each daughter cell. Screenshot from simulation step 1803. 1 simulation step equals 1 hour. (**B’’**) Plot of cell age (hours elapsed since last cell division) in simulation step 1000 showing a gradient of ages with the oldest cells in the virtual embryonic retina. (**C’**) In the inducer growth mode, the modelled tissue signals upstream to drive growth of other tissues in the organ. (**C’’**) Screenshot of a simulation in the inducer growth mode. (**C’’’**) Sample of clones in (C’’) with colors denoting the ratio of full perimeter by bounding rectangle perimeter, a metric for shape complexity. (**D’**) In the responder growth mode, control of the growth of the modelled tissue is downstream of an external signal. (**D’’**) Screenshot of the responder growth mode. (**D’’’**) Sample of clones in (D’’) showing diversity in shapes as evaluated by the same shape metric as in (C’’’). Value range as in (C’’’).

*In vivo*, the spatial extent of the CMZ stem cell domain is believed to be defined by cues such as nearby blood vessels (Wan *et al.*, 2016; Tang *et al.*, 2017). Therefore, we defined the virtual stem cell domain with a fixed size of 25 μm, *i.e.* 5 rows of cells, reflecting the endogenous scale of the Rx2-expressing CMZ domain (Reinhardt *et al.*, 2015; Wan *et al.*, 2016; Tang *et al.*, 2017). *In vivo*, NR stem cells divide predominantly asymmetrically, but also undergo symmetric divisions (Centanin *et al.*, 2014). The rates of asymmetric and symmetric divisions are unknown; likewise, it is unknown whether these rates are deterministically defined or an emergent property of an underlying stochastic system. Since stochastic cell divisions successfully describe the proliferation of committed retinal progenitor cells in larval zebrafish (Wan *et al.*, 2016), we used a simple stochastic mechanism for our initial model. Virtual stem cells divide with a fixed probability of 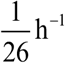, *i.e.* on average once every 26 hours. Intervals between subsequent cell divisions must fulfill a minimum cell cycle length, assumed to be 24 hours. All divisions are symmetric, resulting in two stem cells; cells differentiate and stop cycling when they exit the virtual CMZ after being pushed out by cellular crowding.

To prevent physically implausible cell crowding, cell-center based models include a density-dependent inhibition of cell division (Pathmanathan *et al.*, 2009; Sütterlin *et al.*, 2017). In our model, inhibition occurs in cells whose average overlap with all neighbors exceeds a fraction of the cell’s diameter given by the model parameter *δ*_ol_threshold_ (Figure 2 Supplement 1; Supplementary Text, section 2.4). Based on *in vivo* observations (Lyall, 1957; Johns, 1977; Ohki and Aoki, 1985), the growing virtual eye gradually moves cells apart as it expands, thus decreasing cell density (Figure 2 Supplement 2; Supplementary Text section 2.2). Continuous proliferation in the CMZ counteracts this decrease *in vivo* (Johns, 1977; Johns and Easter, 1977); likewise, the ever-increasing virtual cell population optimally fills the hemisphere at all times (Supplementary Movie 1). Our model distills the complexity of the system to its minimal components and replicates the exquisite spatio-temporal growth order observed *in vivo* (Figure 2 B’, B’’).

### Fundamental feedback modes of organ and cell growth impact on clonal patterns

In a complex organ, feedback mechanisms must coordinate growth of all constituent tissues. Conceptually, this feedback can be wired in two fundamental ways: Either the tissue of interest acts upstream to induce growth of other tissues (Figure 2 C’; “inducer growth mode”), or, vice versa, the tissue of interest lies downstream of growth cues from another tissue in the organ (Figure 2 D’; “responder growth mode”). Possible biological mechanisms for these growth modes could be mechanical, biochemical, or a combination of both. For example, in the inducer growth mode cells could instruct organ growth by modifying the extracellular matrix or by paracrine signalling. These stimuli instruct tissues with the responder growth mode to grow, *e.g.* by alleviating contact inhibition or by providing permissive proliferation signals. In an organ composed of multiple tissues, one tissue may be the driver for growth, while the rest follows.

We examined how these two conceptual growth modes affected clones in the simulation. In our implementation of the inducer growth mode, an increase in cell number induces growth of the virtual eye’s radius (Supplementary Equation 5). Implicit in this growth mode is the assumption that cell division is not inhibited by the degree of cell crowding normally present in the tissue (otherwise the organ would never grow). Therefore, we set the tolerated overlap threshold *δ*_ol_threshold_ = 0.4, a value which we empirically determined to minimize cell division inhibition while preventing physically implausible crowding.

In the responder growth mode, we let the radius grow linearly over time (Supplementary Equation 6). In this growth mode, cells must stop dividing until they receive an external stimulus. We take advantage of the pre-existing local density sensing to implement a physical stimulus akin to contact inhibition. Thus, we set the tolerated overlap threshold *δ*_ol_threshold_ = 0.2 to maximize cell division inhibition at homeostatic density. As growth of the hemisphere decreases cell density, cells dynamically respond to growth of the radius by resuming divisions.

In short, the growth modes in our simulation differ only in: 1) the growth equation for the radius of the hemisphere, 2) the value of the threshold parameter *δ*_ol_threshold_ where local cell density inhibits cell divisions (Supplementary Text, sections 2.3 and 2.4)

We obtained virtual ArCoS regardless of growth mode (Figure 2 C’’, D’’). Strikingly, the growth mode strongly impacted on the shape of ArCoS. Clones in the inducer growth mode formed well-confined stripes with low variation in shape (Figure 2 C’’’). In the responder growth mode, the virtual clones intermingled and frequently broke up into smaller clusters (Figure 2 D’’’). The growth modes impacted on variability in cell division timing. In the responder growth mode, local competition for space increased cell division stochasticity, leading to more diverse clone shapes and clone fragmentation. Thus, the model predicted distinct levels of stochasticity, which, in retinal tissues, resulted in different clonal patterns for inducer or responder growth modes.

### NR stem cells have less stochastic cell division timing compared to the RPE

Due to the spatio-temporal order in the retina, the shape of clones is a readout for proliferation properties of CMZ cells. In the extreme case of no stochastic variation in cell division timing, each clone forms a continuous, unbranching stripe (Figure 3 B, left). In the opposite highly stochastic case, clones frequently branch or merge into polyclones, as well as fragment into several small patches (Figure 3 B, right). Thus, with increasing stochasticity in cell division timing, we expect an increasing incidence of clone branching and fragmentation.

**Figure 3.**
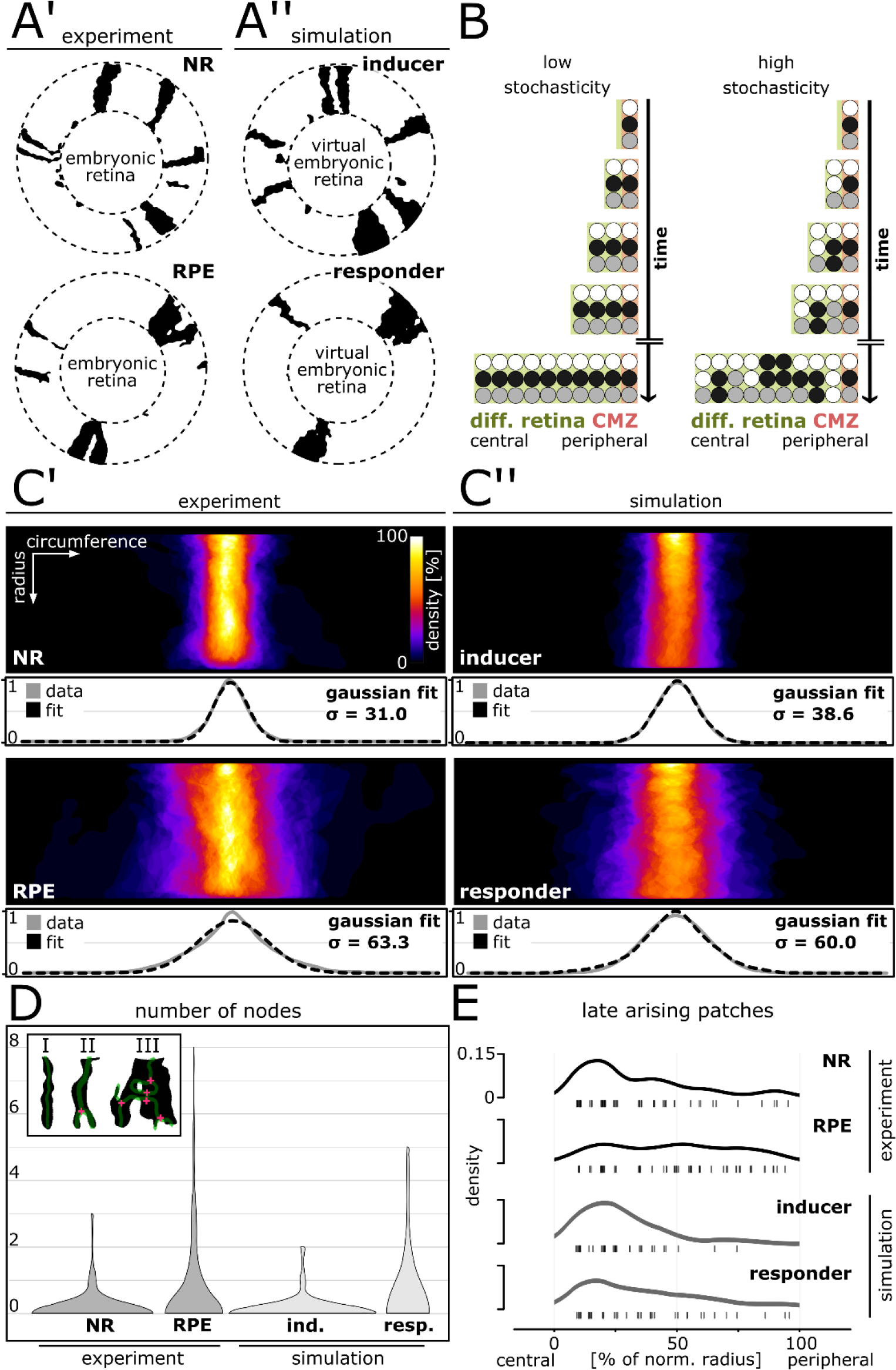
Cell division stochasticity is lower in NR and inducer growth mode, higher in RPE and responder growth mode. (**A’-A’’**) Proximal view of segmented patches in adult NR and RPE and simulated patches in inducer and responder growth mode. The central (virtual) embryonic retina was excluded from analysis. (**B**) Different degrees of stochasticity in cell division timing affect the clone pattern. (**C’-C’’**) Upper panels: Superposition of labelled patches in the NR (n=156 patches from 7 retinae), RPE (n=142 patches from 10 retinae), inducer growth mode (n=145 patches from 5 simulations), and responder growth mode (n=107 patches from 5 simulations). The radius was normalized to the same length in all samples. Lower panels: Gaussian fits of normalized pixel intensity profiles projected along the vertical axis. σ - Standard deviation of fit. (**D**) Distribution of number of nodes of skeletonized patches. Inset: Examples of patches without nodes (I), with only 1 node (II), or with multiple nodes (III). (**E**) Rug plot showing number of patches that are not connected to the embryonic retina “late arising patches”) at the respective positions along the normalized radius. NR (n = 54 late patches) and inducer growth mode (n = 35 late patches) display a marked peak in the central portion, while RPE (n = 56 late patches) and responder growth mode (n = 37 late patches) have a more even distribution.

We compared simulated clones of the inducer and responder growth modes to clones in the NR and RPE (Figure 3 A’, A’’). We circumvented biases associated with fusion and fragmentation of clones by analyzing “patches”, *i.e.* contiguous domains of segmented pixels. A patch may entail a (sub-)clone, or multiple clones (*i.e.* a polyclone) (Figure 3 Supplement 1). To assay the degree of stochasticity in our experimental and simulated data, we quantified patch shape variability, branching, and fragmentation.

We unrolled the retina with a coordinate transform, then aligned and superimposed all patches (Figure 3 C’, C’’). Confirming our previous qualitative observations, NR patches formed a narrow stripe, while RPE patches showed much greater variability (Figure 3 C’). In striking agreement to the experimental data, simulated patches in the inducer growth mode had low variability, while patches in the responder growth mode spread widely (Figure 3 C’’). Patches in the NR and in the inducer growth mode were overwhelmingly stripe-like with no branch points (Figure 3 D; inset I). In contrast, patches in the RPE and in the responder growth mode frequently bifurcated or merged, creating branching shapes with inclusions indicative of clone intermingling (Figure 3 D; inset III).

Not all patches were contiguous with the embryonic retina. Such “late arising patches” result if a cell divides intermittently with periods of dormancy, leaving clone fragments behind (Figure 3 B, highly stochastic scenario). We plotted the number of late arising patches against their central-most position in the normalized post-embryonic retinal radius (Figure 3 E). In the RPE, and to a lesser degree in the responder growth mode, late patches spread evenly across the post-embryonic retinal radius. Such an even distribution of late patches indicated frequent fragmentation throughout the life of the animal as predicted for the highly stochastic scenario. In contrast, in the NR and in the inducer growth mode, late patches clustered in the central post-embryonic retina and waned thereafter. Thus clone fragments were not equally distributed, consistent with lower levels of stochasticity and a majority of continuous stripe-like clones.

Together, these data show that NR and RPE have different degrees of stochastic variability in cell division timing. The NR displays lower stochasticity consistent with the simulated inducer growth mode, while the RPE shows higher levels of stochasticity that may even exceed what we modelled with the responder growth mode. Thus, our data support a model where NR and RPE concertedly expand relying on different growth modes, which manifest in differently shaped ArCoS. The RPE patch pattern indicates a responder growth mode characterized by higher stochasticity driven by instructive signals of the NR as an inducer tissue with lower stochasticity.

### Stem- and progenitor cell domains are an emergent property of the system

Both the NR and simulations displayed a cluster of late patches in the central post-embryonic retina (Figure 3 E), suggesting that beyond fragmentation an additional stochastic process took place after clonal labelling. The region at the border to the embryonic retina, the “induction ring”, marks the original position of the CMZ at the timepoint of Cre-mediated recombination (Figure 4 D, bottom). To investigate how a central peak in late patches related to stem cell dynamics in the induction ring, we turned to the simulation.

**Figure 4.**
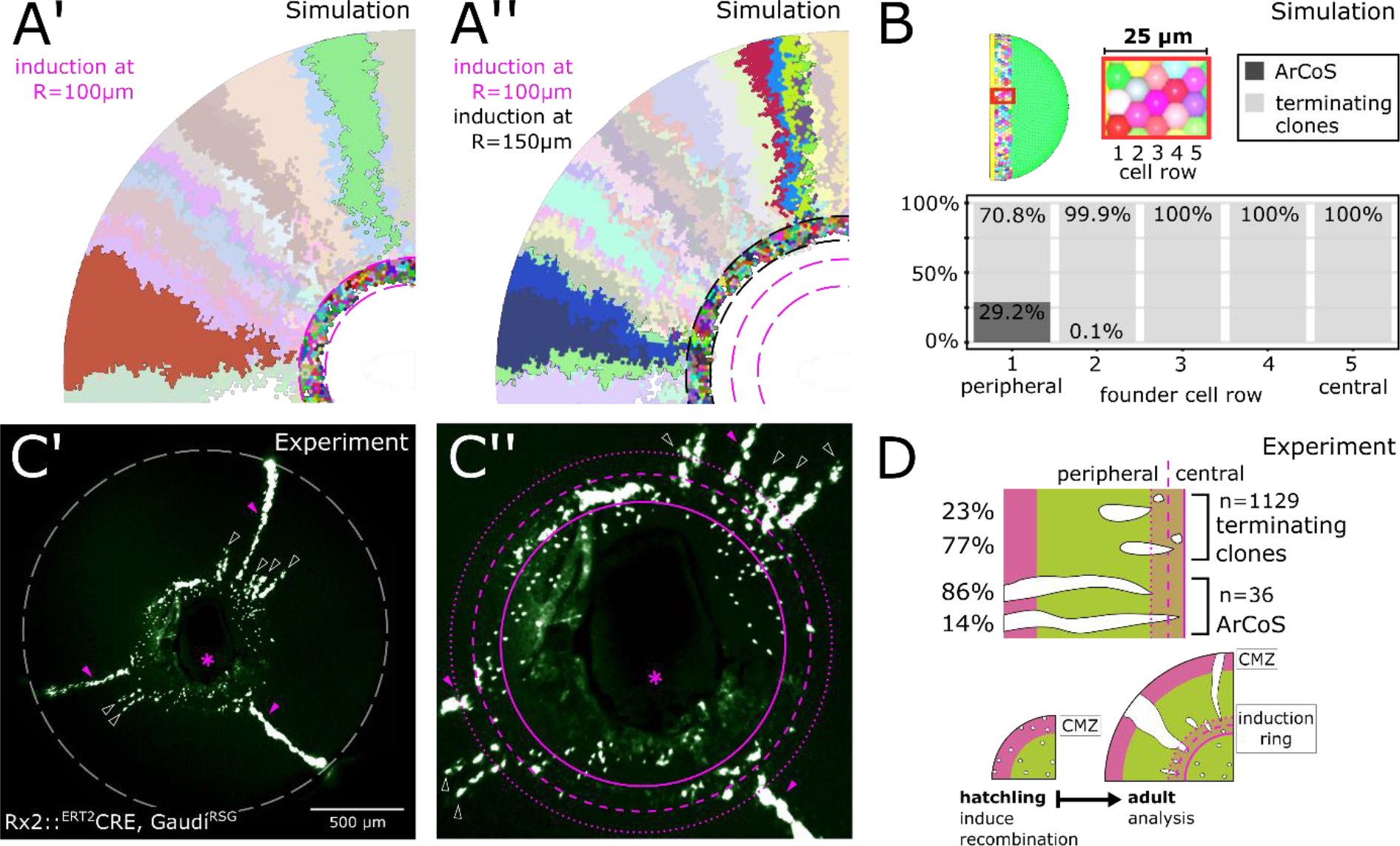
The majority of stem cells differentiates due to cell competition. (**A’**) Detail of inducer growth mode simulation where clone label was initiated at a radius of R=100 μm. Small clusters lie centrally, while virtual ArCoS start peripherally. Two virtual ArCoS are highlighted. Pink dashed lines encircle virtual induction ring. (**A’’**) Same simulation as in (A’), but with clone label initiated at R=150 μm. The second wave of clonal label leads to a renewed occurrence of small clusters. Two polyclonal patches are highlighted, which correspond to subclones of the highlighted clones in (A’). (**B**) The majority of virtual ArCoS in simulations derives from stem cells located in the two most peripheral rows of the virtual CMZ. (**C’**) Proximal view of NR clones. (**C’’**) Magnification of central retina from (C’) showing induction ring (encircled by dashed pink lines of different size subdividing it into peripheral and central domains). Maximum projection of confocal stack of GFP signal in false colors; rotated to place optic nerve exit (pink asterisk) ventrally. Retinal edge marked by white dashed circle; dashed pink lines encircle and subdivide induction ring; pink arrowheads mark ArCoS, clear arrowheads mark terminating clones. (**D**) The majority, but not all, ArCoS arise from stem cells at the very periphery of the CMZ. Terminating clones arise both from peripheral as well as more centrally-located cells in the induction ring. n=36 ArCoS and n=1129 terminating clones were counted in n=20 retinae.

Surprisingly, the virtual induction ring contained many few-cell clones unrelated to any ArCoS (Figure 4 A’, encircled by pink dashed lines). In these clones, all stem cells left the niche and thus differentiated “terminating clones”). Nested inductions showed that sister stem cells within one clone segregated into subclones (Figure 4 A’-A’’, highlighted ArCoS). However, only some of these subclones generated virtual ArCoS. Again, terminating clones clustered in the virtual induction ring (Figure 4 A’’, encircled by black dashed lines), demonstrating that the pattern repeated itself regardless of the timepoint of virtual induction. Therefore, since central positions were occupied by short terminating clones, many stripe-like patches necessarily began in more peripheral positions, explaining the peak in late arising patches.

In our model, all proliferative cells were equipotent stem cells. Nevertheless, a subset of these virtual stem cells proliferated only a few times before terminally differentiating, akin to commited progenitor cells. Notably, the overwhelming majority of virtual ArCoS emerged from the periphery of the induction ring (Figure 4 A’-A’’), as confirmed by tracing back the original position of the founder stem cells, while centrally located cells formed exclusively terminating clones (Figure 4 B). Strikingly, only a minority of stem cells formed ArCoS, while the vast majority formed terminating clones (Figure 4 B).

Together, these data show that the virtual niche subdivided into a peripheral stem- and a central progenitor domain. Importantly, this subdivision was not imposed onto the simulation, but emerged dynamically. The central-most cells were poised to differentiate by stochastically being pushed out of the niche by divisions of their more peripheral neighbors. This neutral competition occurred continuously, as demonstrated by nested virtual inductions (Figure 4 A’-A’’). Thus, the spatial segregation of stem- and progenitor domains was an emergent property of the system.

### Experimental clones follow a spatially biased stochastic drift

Our simulations uncovered a role of stochastic drift in the niche, and lead us to the following two predictions: First, a large proportion of stem cells is lost by neutral competition and forms terminating clones. Thus, ArCoS should be a minority among labelled clones. Second, there is a spatial bias in this drift: The majority of ArCoS will derive from peripheral cells but some will derive from more central positions. Similarly, the majority of terminating clones will derive from central positions, but some will derive from peripheral positions.

To address these predictions experimentally, we again labelled NR stem cells in hatchlings using the Rx2::^ERT2^Cre, Gaudí^RSG^ line (Centanin *et al.*, 2014; Reinhardt *et al.*, 2015), which when recombined results in a nuclear GFP signal, and analyzed the eyes at adult stage. Few-cell clusters in the induction ring vastly outnumbered ArCoS, showing that terminating clones were the most common type of clone (n=1129 terminating clones in 20 retinae; Figure 4 C’-C’’, Figure 4 Supplement 1 A-B). A small fraction of terminating clones extended into the post-embryonic retina (Figure 4 C’-C’’, black arrowheads). ArCoS, which by definition always reach the retinal margin, were less frequent (Figure 4 C’-C’’, pink arrowheads; n = 36 ArCoS in 20 retinae). Thus, Rx2-expressing cells in the CMZ include cells that proliferate indefinitely as well as cells that proliferate only a few times before differentiating. The preponderance of terminating clones shows that ArCoS-forming cells are a minority, in line with our first prediction.

To address the spatially biased stochastic drift, we examined at which position in the induction ring clones derived from. We recorded the proportions of ArCoS and terminating clones arising from the peripheral and central induction ring (Figure 4 D). Among terminating clones, 77.3% started in central positions, while 22.7% were exclusively located in the peripheral induction ring or in the post-embryonic retina. Thus, a subset of terminating clones derived from the periphery of the stem cell domain of the CMZ, indicating that some stem cells drifted into a progenitor-like state. The majority of ArCoS (86.1%) started in the periphery, but 13.9% derived from a central position, suggesting that some cells located in the central progenitor domain of the CMZ drifted into a lifelong stem cell fate. Together, these data support a model of stochastic drift with a peripheral-stem and central-progenitor bias that is conditioned by the physical topology of the niche.

### NR stem cells undergo radial divisions at the rate predicted by shape regulation

Despite stochastic competition among stem cells (Figure 4 C’-D), NR ArCoS formed stripes narrower than in the simulation (Figure 3 C’-C’’). In simulations, the division axis was not oriented (“random division axis”). The thin clonal stripes suggested that NR stem cells had a preferential axis of division along the radial (central-peripheral) coordinate, while circumferential divisions occurred with lower frequency than expected for a random division axis orientation.

We wondered whether NR stem cell division orientation could relate to shaping the organ. An inducer growth mode does not necessarily imply regulation of organ shape. To use an analogy, a mass of dough grows from within (similar to the inducer growth mode), but its shape can be imposed externally by a mold (*i.e.* the dough does not affect shape regulation). In the NR, the shape could plausibly be imposed externally by any of the surrounding tissues, and in this case, it would have no role in organ shape regulation (Figure 5 A). As the space available for cells is imposed externally, any orientation of division axes is theoretically possible; after division cells will locally shift to optimally fill space. In an alternative scenario, organ shape could be regulated by oriented cell divisions of CMZ stem cells (Figure 5 B). In this scenario, a precise orientation of division axes is necessary.

**Figure 5.**
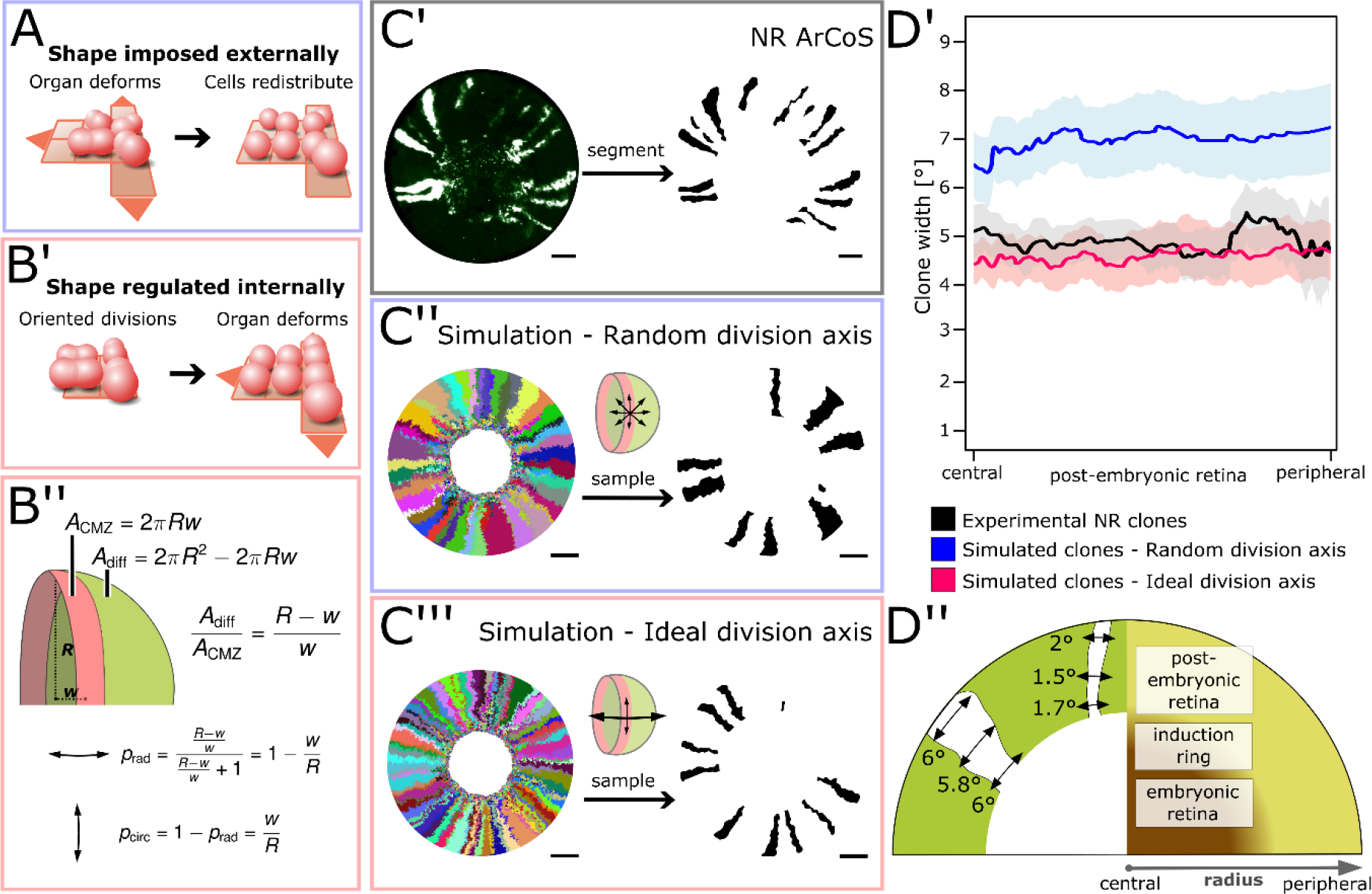
NR stem cells undergo predominant radial divisions as predicted for a shape-giving function. (**A**) If organ shape is imposed externally, then cells in the tissue will distribute to fill the available space. Regardless of cell division axes, organ geometry will lead to a directional growth in stripes. (**B’**) If organ shape is regulated by cell division axes, then oriented divisions are required. (**B’’**) If the NR regulates shape through cell divisions, then more divisions along the radial axis are needed to maintain hemispherical geometry. (**C’-C’’’**) Examples of experimental and simulated data. For simulations, the full clone population and a random sample are shown. The initial model label was induced at R=150 μm to match the experimental induction radius. Scale bars: 200 μm. (**D’**) Mean clone width (solid lines) and 95% confidence intervals (shaded) plotted along the post-embryonic retinal radius. Experimental data: n = 99 ArCoS across 7 retinae. Simulation, random division axis: n = 102 ArCoS from 5 simulations; ideal division axis: n = 133 ArCoS from 5 simulations. (**D’’**) Schematic of radial compartments of the NR and measurements of clone angular width in proximal view.

We calculated the ideal proportion of circumferential and radial divisions required to maintain hemispherical geometry. We assumed two principal axes of division, and that each new cell contributed either to the area of the CMZ or to the rest of the eye (Figure 5 B’). Circumferential divisions (two daughter cells stay in the CMZ) must be balanced by radial divisions (one daughter cell is poised to leave the niche and differentiate). A hemispherical eye of radius *R* has the area

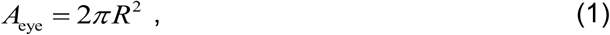

while the CMZ forms a band of width *w* at the base of the eye with area

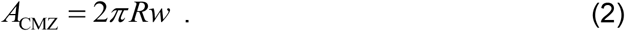

Thus, we obtain an ideal ratio of circumferential to radial divisions of

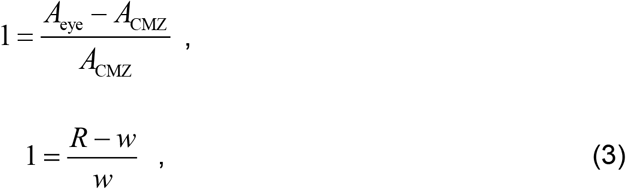

*i.e.* for every 1 circumferential division, there must be 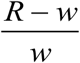 radial divisions. Since *R ≫ w*, this means that radial divisions must be more frequent than circumferential divisions, and the frequency of radial divisions increases as the radius of the retina grows.

When division axes perfectly match the ratio in Equation 3, the simulation becomes the limiting case of shape regulation where the hemispherical shape is always maintained. Thus, we modelled how the “ideal division axis” ratio given by Equation 3 affected the properties of simulated ArCoS and compared this to experimental data as well as simulations with random division axis (Figure 5 C’-C’’’).

To quantify circumferential stem cell divisions in experimental and simulated data, we took advantage of the exquisite temporal order of NR growth to measure ArCoS width – a proxy for circumferential stem cell divisions (Figure 5 D’-D’’). To only include lifelong stem cells, we focused our analysis on the post-embryonic retina and excluded the central portion including the induction ring.

Experimental ArCoS width averaged to 4.87° (Figure 5 D black graph; n = 99 ArCoS across 7 retinae). In contrast to experimental data, ArCoS width in simulations with random division axis averaged to 7.28° (Figure 5 D blue graph; n = 102 clones from 5 simulation runs). In simulations with ideal division axis, ArCoS width closely matched experimental data, averaging at 4.54° (Figure 5 D, red graph; n = 133 clones from 5 simulation runs). These data show that NR stem cell divisions were not completely random, but instead were preferentially oriented along the central-peripheral axis. Moreover NR stem cells underwent radial and circumferential divisions at a rate consistent with a role in organ shape regulation.

### Local biases in ventral NR stem cell divisions influence retinal topology

We observed that in the retina of the surface-dwelling medaka, the position of the embryonic retina was not centered, but instead was shifted ventrally (Figure 6 A). The post-embryonic retina was longer dorsally than ventrally; the ratio of dorsal to ventral length had a mean of 1.42 and a standard deviation of 0.29 (n = 10 retinae). The embryonic retina covered the entire retinal surface at induction (Figure 6 B). Equal growth around the circumference should maintain the embryonic retina in the center. The ventral-ward shift indicated that along the CMZ circumference, ventral stem cells had different division parameters.

**Figure 6.**
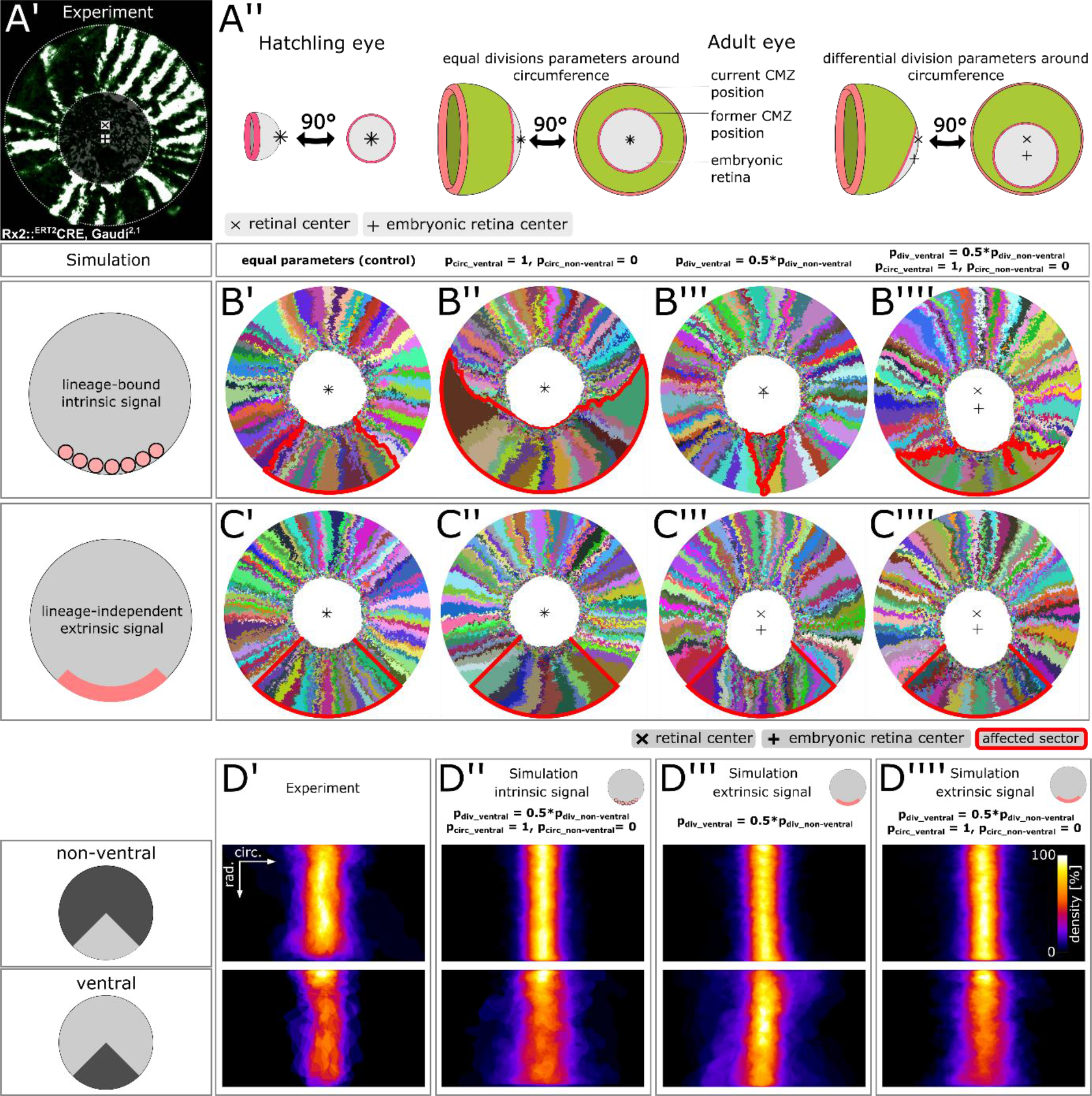
Stem cells in the ventral CMZ have different proliferation parameters. (**A’**) Proximal view of NR clones highlighting the discrepancy between retinal center and embryonic retinal center. Depicted sample is the same as in Figure 1 D. (**A’’**) A differential proliferative behavior along the CMZ circumference can explain the shift in position of the embryonic retina. (**B’-B’’’’**) Simulations where lineages whose embryonic origin is in the ventral sector inherit a signal that leads to different proliferation parameters. A shift occurred when ventral lineages had both lower division probability and circumferential divisions. Clones originating in ventral embryonic CMZ are outlined in red. (**C’-C’’’’**) Simulations where all cells in a ventral 90° sector exhibit different proliferation parameters regardless of lineage relationships. A shift occurred in conditions with slower proliferation as well as slower proliferation combined with circumferential division axis bias. (**D’-D’’’’**) Superposition of non-ventral and ventral patches of experimental data as well as the three simulated conditions that display a ventral shift of the embryonic retina.

We probed the feasibility of different scenarios in generating a ventral shift in an *in silico* screen. First, we discerned two ways for stem cells in the ventral domain (defined as a 90° sector; Figure 6 Supplement 2) to select a different division behavior: Either a lineage-bound intrinsic signal (*e.g.* epigenetic imprinting), or a lineage-independent extrinsic signal (*e.g.* a local diffusible molecule). Second, we altered two cell division parameters: The probability of division, which affects how fast cells divide on average, and the preferential axis of cell division, which we varied between circumferentially-biased and radially-biased.

In control simulations where all cells behaved equally, the embryonic retina stayed centered (Figure 6 B’, C’). For a lineage-bound intrinsic signal, a circumferential division axis bias lead to massive enlargement of ventral lineages at the expense of adjacent clones without affecting the embryonic retina (Figure 6 B’’). Reducing proliferation probability resulted in termination of ventral lineages, as adjacent clones displaced them from the virtual niche (Figure 6 B’’’). An intrinsic signal resulted in a ventral shift only if circumferential division axis bias was combined with lower proliferation probability (Figure 6 B’’’’). In these simulations, circumferential divisions allowed ventral lineages to physically occupy niche positions (preventing their displacement) while lower proliferation reduced pressure on cells of the embryonic retina, allowing a ventral shift. In the scenario of a lineage-independent extrinsic signal, two conditions resulted in a ventral shift of the embryonic retina: Both lower division probability (Figure 6 C’’’) and the combination of lower division probability with circumferential division axis bias (Figure 6 C’’’’).

To identify which scenario was most plausible, we analyzed patches in the ventral and non-ventral sectors. Both in experiments and all three simulated conditions, patch shape in the non-ventral sector was similar (Figure 6 D’-D’’’’, top panels). Importantly, although there was a tendency for ventral clones to terminate more often, experimental NR patches did not differ substantially in width between non-ventral and ventral sectors (Figure 6 A, D’). In contrast, this latter criterion was violated by two of the three simulated scenarios (Figure 6 D’’-D’’’’, Figure 6 Supplement 1 A-C).

When an intrinsic signal instructed both lower proliferation and a circumferential bias, ventral ArCoS started narrow but then broadened (Figure 6 E’’) and interdigitated laterally (Figure 6 Supplement 1 A, white arrowheads), unlike the very uniform stripes in the experimental data. When an extrinsic signal reduced proliferation probability, the majority of ventral ArCoS formed very narrow stripes, but at the border to the non-ventral domain ArCoS were broad and curved (Figure 6 Supplement 1 B, white arrowheads). Again, this resulted in more shape variation (Figure 6 E’’’). Ventral clones were highly similar to each other and the rest of the retina only in the condition where an extrinsic signal both lowered proliferation and imparted a circumferential division axis bias (Figure 6 E’’’’, Figure 6 Supplement 1 C). This condition also displayed a similar trend for terminating ventral clones (higher patch density in central position) and thus the overall greatest congruence with the experimental data.

In conclusion, ventral NR stem cells have a different behavior than elsewhere along the circumference, leading to a ventral-ward shift of the embryonic retina. The simulations suggest that this different behavior consists of concurrent modulation of both proliferation rate and division axes by an extrinsic signal in the ventral CMZ.

## Discussion

### The NR drives growth upstream of the RPE

The coordinated growth of multiple independent tissues is a ubiquitous process in biology. In this work, we used the post-embryonic growth of NR and RPE in the eye of medaka as a model system of coordination in an organ where both growth and shape must be precisely regulated.

Eye size in fish scales to the body size (Lyall, 1957; Johns and Easter, 1977). Body size, and thus eye growth rates greatly vary among individuals and environmental factors (Johns, 1981). This natural malleability implies that feedback coupling plays a dominant role rather than the precise parametrization of each tissue growth rate. Strikingly, our simulations showed that inducer and responder growth modes impacted on stochasticity in cell divisions, ultimately resulting in distinct clonal patterns that reproduced the experimentally observed differences between NR and RPE.

RPE cells divided with high stochasticity, indicative of periods of long dormancy where they waited for proliferative cues. NR cells displayed lower stochasticity, supporting an upstream role in regulating growth (Figure 7 A). Although our implementation of the responder growth mode used a mechanical stimulus (local cell density), a biochemical stimulus could equally well represent the system.

**Figure 7.**
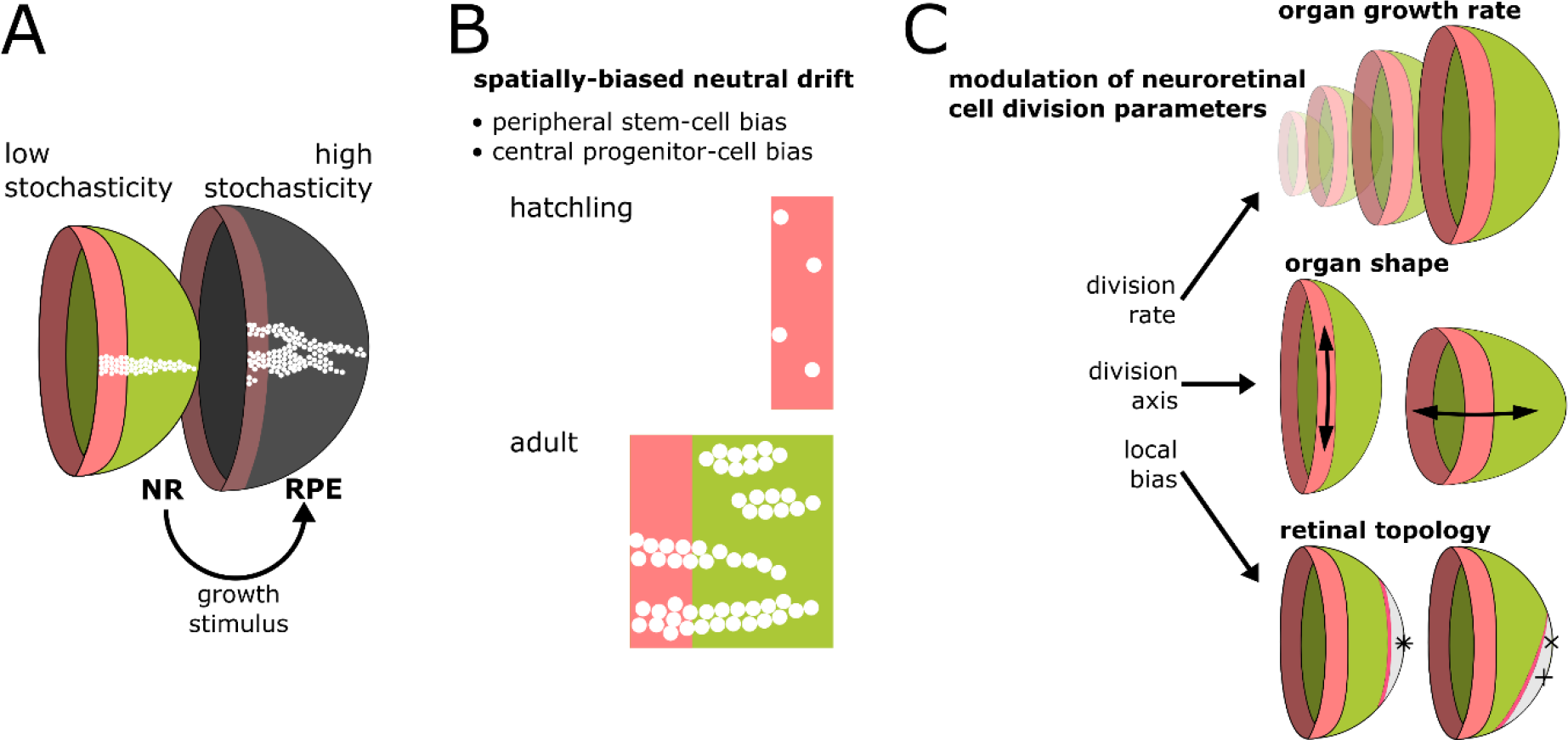
Summary of results and proposed model of CMZ dynamics. (**A**) Growth coordination of NR and RPE is achieved by the NR providing instructive stimuli that modulate proliferation of RPE stem cells. As a result of the different growth strategies, proliferation stochasticity is elevated in the RPE and lowered in the NR. (**B**) A base level of stochasticity persists in the NR, such that individual stem cells may differentiate and some multipotent progenitor cells drift to a stem cell fate according to a spatially biased neutral drift model. Thus, stem cells and multipotent progenitor cells have identical potency. (**C**) Schematic summary of findings and proposed model, where different NR cell proliferation parameters affect both global and local retinal properties.

Our model highlights an underappreciated mechanism whereby tissues coordinate by inducer and responder roles. Such division of labor among tissues might apply more generally to multiple organ systems, *e.g.* hair follicle cells in mouse induce the growth of underlying adipose tissue through hedgehog signalling (Zhang *et al.*, 2016). Intriguingly, hedgehog signalling also regulates the NR/RPE boundary in the CMZ of medaka (Reinhardt *et al.*, 2015), suggesting that signals mediating coordination of proliferative cell populations might be conserved.

### Multipotent progenitor cells are stem cells that were outcompeted

The topology of the retinal niche lead to a spatially biased neutral drift where stem and progenitor compartments spontaneously emerged. All virtual cells had equal potency, yet only a fraction realized their full stem cell potential. Peripheral cells had a high chance to become canalized in a stem cell fate, while central cells were more likely to act as progenitor cells with limited proliferation potential (Figure 7 B).

Our experimental data support a spatially biased neutral drift. Fusion of clones may have lead us to overestimate ArCoS deriving from the central domain, which represent progenitors reverting to a stem cell fate. Nevertheless, terminating clones arising from the very periphery of the niche unambiguously demonstrate that some stem cells failed to self-renew throughout the life of the animal. Moreover, our finding that only cells in the first two rows of the CMZ have stem cell potential is consistent with *in vivo* time-lapse data (Wan *et al.*, 2016; Tang *et al.*, 2017). Interestingly, retrograde movement of row 2 cells into row 1 of the CMZ occurs *in vivo* (Wan *et al.*, 2016), which we also observed in our simulations.

CMZ progenitor cells can be subdivided into two populations (Harris and Perron, 1998; Raymond *et al.*, 2006): First, peripheral multipotent progenitors (*i.e.* able to generate all retinal neurons and glia) which differ from stem cells only in their proliferative potential. Second, central progenitors that are restricted both in proliferative and differentiation potential, which likely act as a transit-amplifying zone, both increasing the proliferative output and cross-regulating to produce a full neuronal complement with the correct proportions of cell types (Perez-Saturnino *et al.*, 2018).

Our data support an alternative model that identifies peripheral multipotent progenitors as stem cells that have been outcompeted. All terminating clones we examined were multipotent and spanned all retinal layers (Figure 1 Supplement 2). Thus, as in many other systems (Clevers and Watt, 2018), our work highlights the limitation of strictly defining stem cells as infinitely self-renewing, or *a posteriori* based on their ArCoS-forming capacity.

Importantly, although stochastic competition is most apparent in the early phase after clonal induction, it occurs continuously as demonstrated by late arising patches (Figure 3 E) and nested inductions (Figure 4 A’-A’’). The shift from an “early stochastic” to “late polyclonal” growth observed in other systems (Nguyen *et al.*, 2017) may simply result from clonal growth masking the underlying stochasticity. Due to this stochasticity, it is impossible to tell at any moment with absolute certainty if a given cell will perpetually function as a stem cell.

### Why does the CMZ niche of the retina not drift to monoclonality?

Neutral drift in a finite-sized environment such as adult mammalian tissues must ultimately result in a monoclonal niche (Snippert *et al.*, 2010; Colom and Jones, 2016; Clevers and Watt, 2018). In fish, homeostatic growth expands niches, and thus the number of stem cells increases (Centanin *et al.*, 2011). In principle, niche expansion reduces the impact of random competition on clonal loss, but does not completely abolish it.

Indeed, neutral drift leads to gradual loss of polyclonality in the intestine and muscle of fish (Aghaallaei *et al.*, 2016; Nguyen *et al.*, 2017). Organs may limit monoclonal drift by physically isolating niches (Aghaallaei *et al.*, 2016). In the intestine of both mammals and fish, physical niche isolation results in a polyclonal organ built up of monoclonal units (Snippert *et al.*, 2010; Aghaallaei *et al.*, 2016). In contrast, the CMZ is a physically contiguous niche that nevertheless maintains polyclonality lifelong both in the NR and the RPE (Centanin *et al.*, 2011, 2014). As shown in this work, the retina is not devoid of stochastic competition. Then how does it conserve its polyclonality?

Conceptually, the clonal growth of the retina resembles a population expanding into a new habitat, as studied in the context of evolutionary theory (Hallatschek and Nelson, 2010). Specifically for a radially expanding population, it has been mathematically proven that (assuming pure neutral genetic drift) no single clone will ever take over and clonal sectors perpetually coexist (Hallatschek and Nelson, 2010; Korolev *et al.*, 2012). Growth of the perimeter is faster than circumferential expansion of clones, thus preserving population diversity (Hallatschek and Nelson, 2010). In the NR, central-peripheral divisions further reduce competition (thus increasing niche polyclonality). In summary, the geometry of the CMZ niche prohibits the total loss of polyclonality.

### The NR senses the retinal radius and directs cell divisions to adapt organ shape

Our analysis of NR stem cell divisions implies that cells sense the radius of the eye to regulate organ shape. Across vertebrates, the retina integrates visual input to adapt organ shape to optimize optics, a process called “emmetropization” (Wallman and Winawer, 2004). In chicken, emmetropization is regulated by specialized neurons distributed across the retina that send their axons to the CMZ, implicating the CMZ in regulation of eye shape (Fischer *et al.*, 2008). Visual cues also guide emmetropization in fish (Kröger and Wagner, 1996; Shen *et al.*, 2005; Shen and Sivak, 2007). Eye growth in young fish predominantly occurs by cell addition, while in older fish CMZ proliferation decreases (Johns, 1981) coincident with a decrease in emmetropization plasticity (Shen and Sivak, 2007). Thus, in fish, emmetropization correlates with CMZ proliferation.

Experiments in chicken and zebrafish support the existence of two principal axes of stem cell division, *i.e.* circumferential and central-peripheral (Fischer *et al.*, 2008; Ritchey *et al.*, 2012; Wan *et al.*, 2016). Notably, the predominance of central-peripheral divisions and decreasing frequency over time of circumferential divisions in CMZ stem cells that is predicted by Equation 3 is supported by *in vivo* imaging data (Wan *et al.*, 2016) and previous long-term clonal analyses (Centanin *et al.*, 2014). Altogether, the data support a model where the NR perceives the retinal radius through visual cues, and that cell divisions in the NR contribute to shaping the eye.

### An eye-internal signal directs local proliferation parameters in the CMZ

The retinae of many fishes grow asymmetrically, perhaps to maintain the relative positions of receptive fields of neurons (Johns, 1977, 1981; Easter, 1992). Ecology dictates a distribution of subdomains enriched in specialized circuits and retinal cell subtypes (Zimmermann *et al.*, 2018). Interestingly, in green sunfish, the area that grows slowest displays highest visual acuity (Cameron, 1995). Medaka predominantly gaze upwards in their native shallow rice paddies, and a higher ventral acuity has been presumed based on photoreceptor densities (Nishiwaki *et al.*, 1997). Thus, slower ventral growth may have evolved to match ecological requirements for medaka vision.

Our *in silico* screen identified three scenarios consistent with asymmetric ventral growth. Based on clonal patterns, an extrinsic signal driving lower proliferation and circumferential divisions appears most plausible. Experimental eye re-orientation *in vivo* implied an eye-internal mechanism independent on body axes or visual cues in regulating retinal asymmetry (Cameron, 1996). The origin of this signal and how it scales with the growing eye to always affect a similarly-sized retinal sector remains to be elucidated.

### The CMZ integrates cues to direct eye growth and shape

The retina integrates global cues such as nutrition to scale with body size (Johns and Easter, 1977), local eye-internal cues to generate an asymmetric retinal topology (Cameron, 1996), and external visual cues to adapt the shape of the organ (Kröger and Wagner, 1996; Shen and Sivak, 2007). In chicken and goldfish, visual cues and nutrients feed into the CMZ through growth factor signalling (Boucher and Hitchcock, 1998; Fischer *et al.*, 2008; Ritchey *et al.*, 2012). We propose that NR cells in the CMZ act as a hub to coordinate organ growth; in the eye of fish, this happens at the level of cell proliferation parameters, which affect eye growth, eye shape, and retinal topology (Figure 7 C).

Indeterminate, lifelong growth is a widespread evolutionary strategy (Karkach, 2006). Given the geometrical constraints of the eye with respect to optics, a peripheral proliferative domain is the most parsimonious architecture to ensure that the differentiated neuronal cell mosaic is not disturbed by constant proliferation. Fishes are the largest vertebrate clade, with a huge diversity of eye shapes, such as cylindrical eyes in deep-sea fish (Fernald, 1990). By modulating CMZ proliferation parameters, evolution can adapt whole-organ morphogenesis to perfectly fit to the species’ ecological niche.

## Materials and Methods

### 1. Key resources table

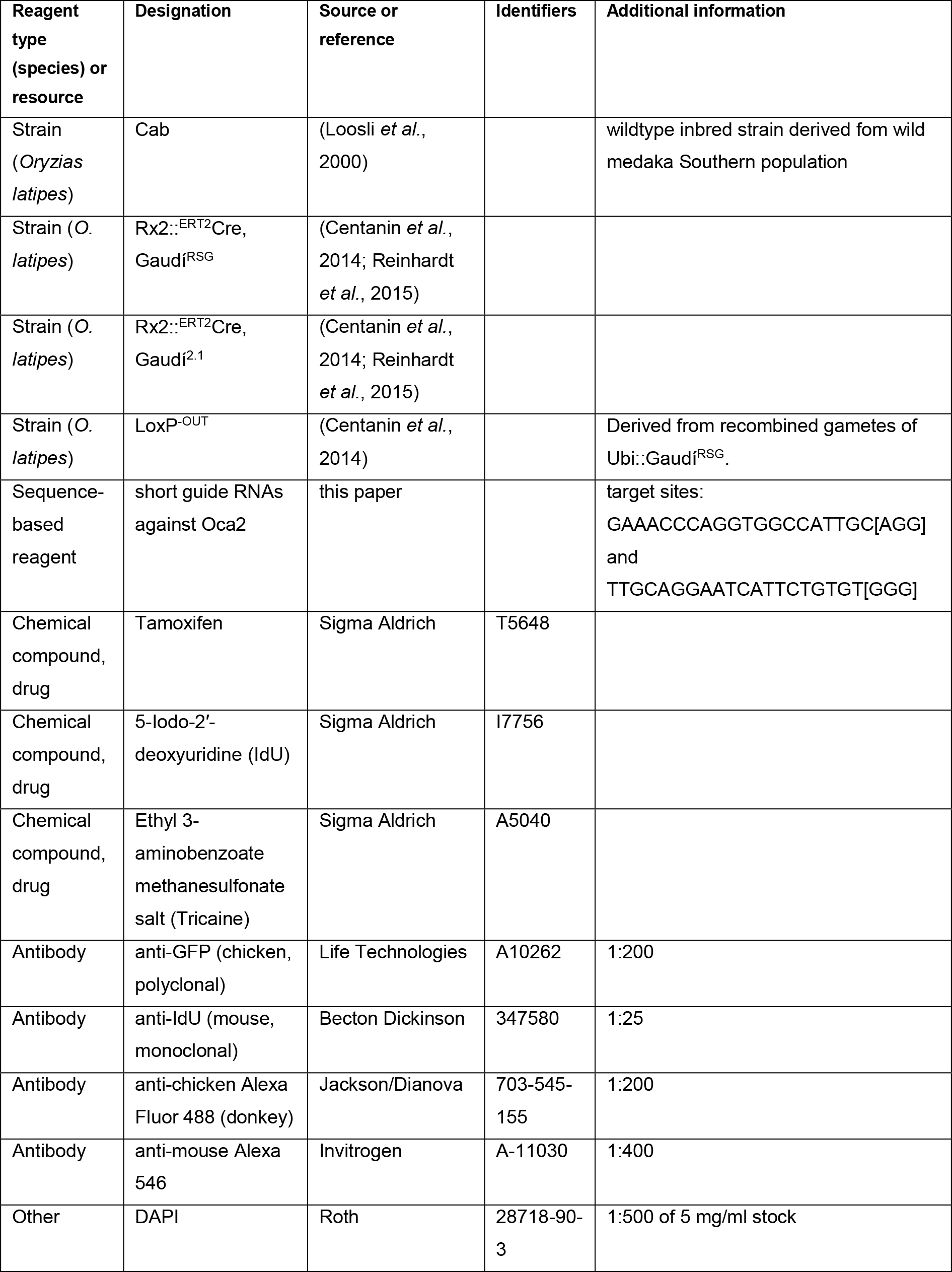

### 2. Experimental methods

#### Animal welfare statement

Medaka (*Oryzias latipes*) fish were bred and maintained as previously established (Loosli *et al.*, 2000). All experimental procedures were performed according to the guidelines of the German animal welfare law and approved by the local government (Tierschutzgesetz §11, Abs. 1, Nr. 1, husbandry permit number AZ 35-9185.64/BH; line generation permit number AZ 35-9185.81/G-145-15).

#### Clonal lineage labelling

ArCoS in the NR were generated as described previously (Centanin *et al.*, 2011, 2014; Reinhardt *et al.*, 2015). Transplantations were from labelled donor cells of the LoxP^OUT^ line to unlabelled wildtype Cab host blastulae. Cre-mediated recombination was performed in hatchlings by induction of the Rx2::^ERT2^Cre, Gaudí lines with 5 μM tamoxifen diluted in fish water for at least 3 hours.

For ArCoS in the RPE, mosaic unpigmented clones were generated using CRISPR/Cas9 by injecting 30 ng/μl each of two short guide RNAs directed against the gene Oca2 in one-cell stage Cab medaka embryos. Oca2 is required to produce melanin pigment (Fukamachi *et al.*, 2004). The sgRNA was designed using CCTop (Stemmer *et al.*, 2015).

#### Treatment with IdU

Fish were bathed in fish water containing concentrations of 2.5 mM IdU as previously described (Centanin *et al.*, 2011).

#### Sample preparation and imaging

Fish were allowed to grow and sacrificed as young adults with an overdose of Tricaine. Whole fish were fixed in 4% formaldehyde in phosphate buffered saline and 0.1% Tween at least once overnight at 4°C while gently shaking. Eyes were dissected, if necessary immunostained, and imaged at a Nikon AZ100 upright stereomicroscope using a 5x dry objective.

#### Immunostaining

To remove melanin pigment, fixed samples were bleached with 0.3% H_2_O_2_ and 0.5% KOH dissolved in phosphate buffered saline and 0.1% Tween (PTW). Samples were permeabilized in acetone for 10 minutes at −20°C. Blocking was performed for at least one hour in a solution of 4% sheep serum, 1% bovine serum albumin (BSA), and 0.1% DMSO, diluted in PTW. Samples were incubated with primary antibodies diluted in 4% sheep serum and 1% BSA in PTW at least once overnight at 4°C with gentle mixing. Secondary antibodies were diluted in 4% sheep serum and 1% BSA in PTW together with DAPI; samples were incubated in secondary antibody solution at least once overnight at 4°C with gentle mixing.

An antigen retrieval step was performed prior to IdU staining. This step consisted of post-fixation in 4% formaldehyde for 1 hour, DNA denaturation with 2M HCl and 0.5% Triton for 45 minutes, and pH recovery for 10 minutes in a 40% borax solution in PTW.

### 3. Data Analysis

#### Experimental clone segmentation

All image processing and analysis was performed using the Fiji distribution of ImageJ (Schindelin *et al.*, 2012). Experimental retinae were selected such that only sparse labelled eyes of comparable size were used for analysis. For NR samples, a maximum intensity projection of confocal stacks was used for segmentation. For RPE samples, a custom script was written to create a focused reconstruction from multiple focal planes based on the hemispherical shape of the whole-mount retina. Briefly, the regions in focus in a stack through a hemispherical object are rings of increasing radii (and a circle in the first plane). The size of these rings was calculated based on the size of the sample. The focused areas were extracted and collated in one composite image.

Labelled clones were segmented by subtracting background noise with a difference of gaussians, and thresholded by the Phansalkar local threshold algorithm as it is implemented in Fiji (Phansalkar *et al.*, 2011). The segmentation was manually curated to eliminate errors.

#### *In silico* clonal lineage labelling

For simulating NR clones, all proliferating cells in the model received a unique ID when the eye radius reached 150 μm. The radius was chosen based on the estimated radius of the NR when genetic recombination was induced *in vivo*. To replicate RPE clones, the virtual labelling experiment began at 100 μm, since mosaic knockout happens at an earlier timepoint in development. The ID is inherited to daughter cells, allowing to reconstruct a lineage at any time during the simulation.

Shape complexity of simulated clones from simulation screenshots was quantified by thresholding individual clones by color, calculating the pixel perimeter, and dividing it by the pixel perimeter of the smallest bounding rectangle enclosing the clone.

For comparison to experimental data, between 8-13% of clones were randomly sampled from the full simulated population; the sample was chosen to produce a sparse label with a comparable number of patches per retina as in the experimental data. Each simulation was sampled twice. The sample of simulated clones was plotted as a 2D projection using a custom Python script; cellular edges were blurred by application of a median filter and shape smoothing plugin in ImageJ.

#### Patch shape analysis

Data analysis on experimental and simulated data was performed using the same automated pipeline in ImageJ, which takes as an input segmented images where the embryonic retina and retinal margin have been previously marked manually. The size of the embryonic retina was estimated based on the induction ring and position of the optic nerve exit, the radius of this estimate was then increased to ensure complete exclusion of all embryonic area. Different sizes of this estimate produced comparable results.

The analysis pipeline treated proximal views of experimental and simulated retinae as polar plots and projected them onto a cartesian coordinate system. After this transform, the width of the image corresponded to the circumference, while the height corresponded to the radius. Radii were normalized to extend from 0% (the border of the embryonic retina) to 100% (the retinal margin). Patch outlines were automatically extracted and superimposed to generate patch density plots. The “plot profile” function in ImageJ was used to extract average pixel intensities along a rectangle spanning the entire image. Gaussian fit was produced in R (R Core Team, 2015).

Skeletonization of patches for node counting was performed using a custom algorithm tailored to the radially oriented retinal lineages: Segmented patches were broken up into radial segments along normalized radial bins ranging from the embryonic to the retinal margin. Each segment was assigned a skeleton element, and these elements were linked in a final step prior to node counting. For each patch, the starting position along normalized radial bins was noted. These data were used to generate the rug plot of late arising patches in R.

#### Quantification of ArCoS and terminating clones

In simulated data, ArCoS were defined as clones that still retained cells in the virtual CMZ at the final simulation step used for analysis, *i.e.* when the virtual retina had attained a radius of R = 800 μm. All other clones counted as terminating clones. The initial position of the founder stem cells for each clone was recorded and assigned to a 5 μm-wide bin corresponding to each of the cell rows in the virtual CMZ.

In experimental data, the position of the induction ring was estimated based on the following criteria: The inner circle was placed such that it enclosed as many 1-cell clones as possible (*i.e.* labelled differentiated cells). The outer circle was placed such that it enclosed all few-cell clusters and crossed all ArCoS. Variation of the position of these two boundaries produced similar results. The induction circle was split in the middle, and the data were segmented as described in “Experimental clone segmentation”. Each clone was assigned to the central-most or peripheral-most ring based on the position of its central-most pixel.

#### Angular width of clones

Both experimental and simulated data were projected onto a rectangular coordinate system as described in the “Patch shape analysis” paragraph. The angular width of clones was measured using a custom script that normalizes the width in pixels to the circumference to obtain the angle. These angular width measurements were exported for analysis and plotting in R. To evaluate only lifelong stem cell clones, the induction ring and small clones that did not extend more than 10% of the radius past the induction ring were excluded from the analysis. Near the retinal margin, the fluorescent signal tapers off due to the retinal curvature and optical limitations of the imaging setup. Thus, the last 5% of the retinal radius were excluded from the analysis. The mean and 95% confidence interval were calculated for each radial position.

## Acknowledgments

We thank the Wittbrodt department, S. Lemke, U. Schwarz, I. Lebovka, E. Kuchen, R. Hodge, and M. González-Gaitán for critical reading of the manuscript. We thank T. Thumberger and M. Stemmer for their help in designing short guide RNAs for CRISPR/Cas9 experiments. We are grateful to A. Saraceno, E. Leist and M. Majewski for fish husbandry. Simulations in this work were performed on the computational resource bwUniCluster funded by the Ministry of Science, Research and Arts and the Universities of the State of Baden-Württemberg, Germany, within the framework program bwHPC. E.T. is recipient of a MSc/PhD fellowship of the Heidelberg International Graduate School for life sciences HBIGS, and of an Add-On Fellowship for Interdisciplinary Science of the Joachim Herz Stiftung. This work was supported by the Research Training Group “Mathematical Modelling for the Quantitative Biosciences” (N.G.), and by the 7th framework program of the European Union (ERC advanced grant GA 294354-ManISteC, J.W.).

The authors declare no competing financial interests. Correspondence and requests for materials should be addressed to E. T. or J. W.

## Author contributions

E.T., B.H., and L.C. performed clonal induction, dissection, and imaging on neural retina samples. E.T. and S.K. performed clonal induction, dissection, and imaging on retinal pigmented epithelium samples. E.T. and B.H. developed quantification tools and analyzed the data. E.T., T.S., N.G., and J.W. conceived the model. T.S. implemented the initial model in EPISIM. E.T. and T.S. extended the model. ET performed and analyzed simulations. E.T., B.H., L.C., and J.W. interpreted and discussed the results. E.T. and J.W. wrote the manuscript. All authors discussed and edited the manuscript.

